# Efficient, high-throughput ligand incorporation into protein microcrystals by on-grid soaking

**DOI:** 10.1101/2020.05.25.115246

**Authors:** Michael W. Martynowycz, Tamir Gonen

## Abstract

A method for soaking ligands into protein microcrystals on TEM grids is presented. Every crystal on the grid is soaked simultaneously using only standard cryoEM vitrification equipment. The method is demonstrated using proteinase K microcrystals soaked with the 5-amino-2,4,6-triodoisophthalic acid (I3C) magic triangle. A soaked microcrystal is milled to a thickness of 200nm using a focused ion-beam, and microcrystal electron diffraction (MicroED) data are collected. A high-resolution structure of the protein with four ligands at high occupancy is determined. Compared to much larger crystals investigated by X-ray crystallography, both the number of ligands bound and their occupancy was higher in MicroED. These results indicate that soaking ligands into microcrystals in this way results in a more efficient uptake than in larger crystals that are typically used in drug discovery pipelines by X-ray crystallography.

## Introduction

Atomic resolution structures are critical to understanding how proteins and small molecules interact. Crystal structures of small molecule pharmaceuticals are also important to determine how drugs are formulated in their delivery (Datta and Grant, 2004). Knowing how the atoms in a molecule are arranged allow for rational design of ligands such as drugs to bind and inhibit or stimulate protein function (Daga et al., 2010; He et al., 2010). Investigation of protein structures with these small molecule ligands bound is an important step in drug discovery and optimization (Blundell, 2017; Kuhn et al., 2002; Tickle et al., 2004). Macromolecular crystallography has been adapted to investigate these ligand-protein interactions by incorporating the ligand into the protein crystal lattice by either soaking ligands into crystals or, more often, by co-crystallizing the protein with the ligand (Beck et al., 2009). Successfully coercing ligands into protein crystals is challenging (McNae et al., 2005). Co-crystallization of the protein with the ligand can change the crystallization condition or result in the ligand forming close contacts between protein molecules in the lattice instead of effectively diffusing into the binding site (Reynolds, 2014). Soaking already formed crystals with ligands is also used (Beck et al., 2009; Lebioda and Zhang, 1992; López-Jaramillo et al., 2002). Crystals soaked with ligands will often crack or dissolve as the additional ligand diffuses into the protein lattice. Cracked or dissolved crystals typically do not diffract well, resulting in either a low-resolution structure or no structure at all. Moreover, soaking ligands into large crystals is difficult because diffusion over long distances is inefficient leading to low occupancy in the crystallographic maps and an ambiguous solution. Therefore, a large challenge in ligand soaking experiments for large macromolecular crystals arises in modeling the weak density that the ligands occupy (Liebschner et al., 2017; Pearce et al., 2017; Pozharski et al., 2013).

Microcrystal electron diffraction (MicroED) is an electron cryo-microscopy (cryoEM) method for determining atomic resolution structures from protein microcrystals (Brent L Nannenga et al., 2014; Brent L. Nannenga et al., 2014; Shi et al., 2013). MicroED has been used to solve novel protein and small molecule pharmaceutical crystal structures (de la Cruz et al., 2017; Gruene et al., 2018; Jones et al., 2018; Rodriguez et al., 2015; Xu et al., 2019). Previous ligand-protein interactions determined using MicroED relied on co-crystallization or incubation of the protein and ligand in crystallization drops prior to applying the crystals to the grid (Clabbers et al., 2020; Purdy et al., 2018; Seidler et al., 2018). In this way, novel co-crystals of the HIV-GAG bevirimat complex were determined (Purdy et al., 2018). The first-generation HIV-I maturation inhibitor bevirimat was found to occupy the six-fold axis of the protein hexamer along the c axis of the unit cell. The density suggested that the drug bound in this position to any of the six protein interfaces, and served as a non-specific inhibitor of the protein capsid, ultimately preventing maturation of the viral capsid. A more recent investigation of protein microcrystals using MicroED was also able to correctly locate the inhibitor acetazolamide (AZM) in the active site of active site of human carbonic anhydrase isoform II (HCA II) after adding AZM to the crystallization drops (Clabbers et al., 2020). Time resolved experiments that expose the protein to a ligand just prior to vitrification for cryoEM imaging were similarly pioneered by Unwin using acetylcholine receptor helical tubes (Unwin, 1995).

We now demonstrate a simple method for efficiently soaking small molecule ligands into pre-formed microcrystals directly on the TEM grid just prior to vitrification. The method involves first applying crystals to a grid similarly to any other cryoEM experiment, back blotting, adding the ligand solution, and back blotting again prior to vitrification by plunging into liquid ethane. This method is demonstrated by on-grid soaking the I3C “magic triangle” ligand into microcrystals of the enzyme proteinase K. On-grid soaking is coupled with cryo-FIB milling which removes any excess material surrounding the crystals thereby increases the quality of data from the selected crystal as inelastic scattering is minimized and the ideal crystal thickness is used. Cryo-FIB milling also allows investigation of ligand soaked protein crystals of any size, that greatly expands the scope of the method (Duyvesteyn et al., 2018; Martynowycz et al., 2019). The data shows that the I3C ligand soaks into the crystals with higher efficiency when compared to using X-ray crystallography with much larger crystals obtained under similar conditions. These results have far-reaching implications in future investigations utilizing MicroED for drug discovery and optimization. The approach may also serve as a platform for soaking experiments of interest in other cryoEM modalities, such as single particle or cryotomography.

## Results

### Soaking the ligand into the crystals

Proteinase K crystals were grown in batch to an average size of 1-20μm. Crystal slurry was applied by a pipette to the carbon side of the grid, and the grid was blotted from the back inside a temperature and humidity controlled blotting chamber. In this way, the liquid flows through the holes of the holey carbon film and the crystals remain on the grid. The grid essentially acts as a filter for the crystals. To the grid of blotted crystals, I3C solution was added and allowed to incubate. This step is mimetic of soaking a looped crystal into a ligand solution as typically done in X-ray crystallography (Beck et al., 2009; McNae et al., 2005). Following incubation, the grid was blotted from the back again, and immediately plunged into liquid ethane (Figure 1). The grids were then transferred to either a Thermo-Fisher Aquilos dual beam FIB/SEM or a Thermo-Fisher Talos Arctica TEM for inspection (Figure 1).

**Figure 1.**
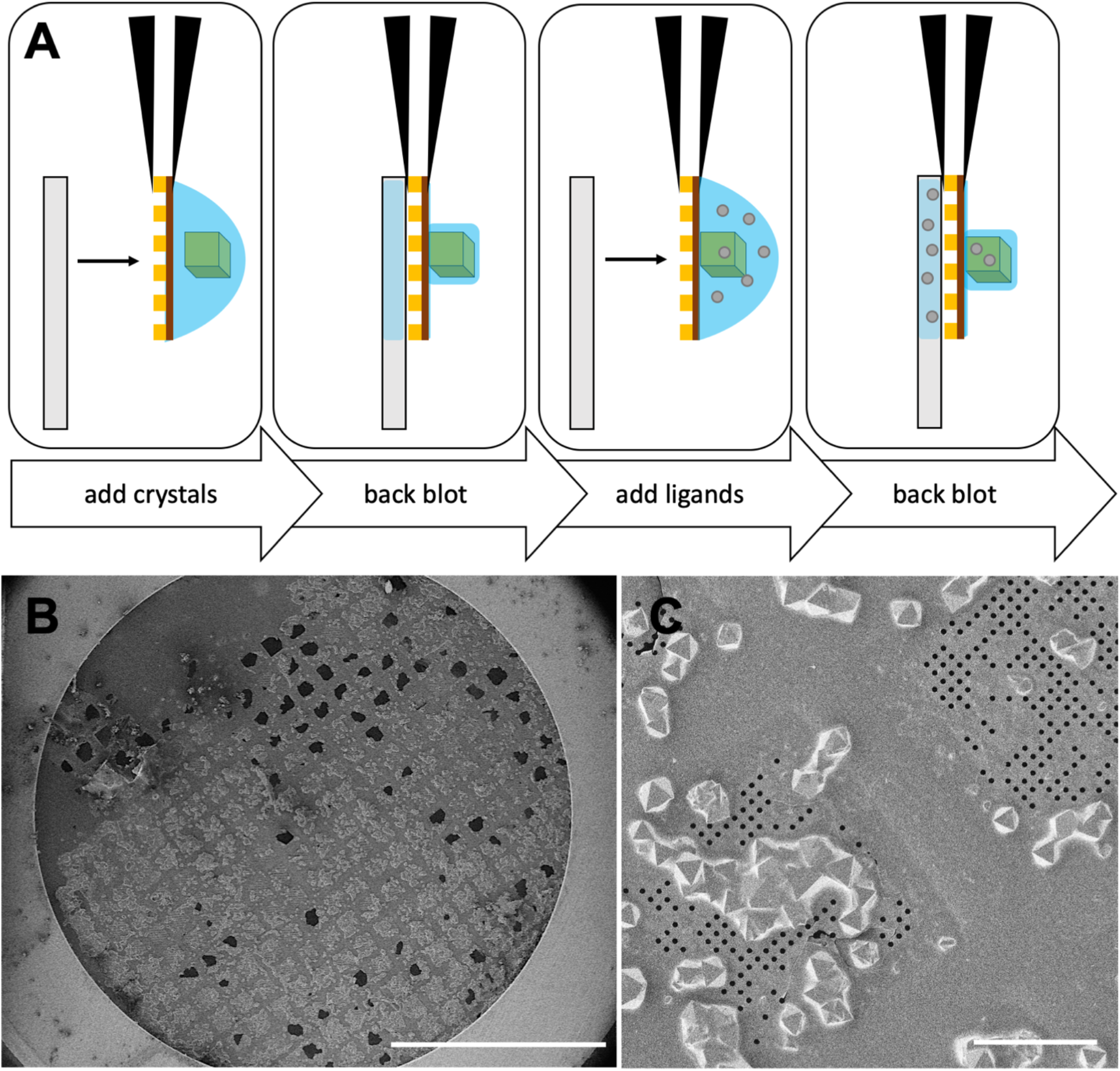
On-grid soaking of ligands into protein microcrystals. (A) Schematic cartoon of the on-grid soaking procedure. From left to right: protein solution is applied to the carbon side of a glow-discharged holey carbon film, the grid is blotted from the back in high humidity for ~20s, the ligand solution is applied to the same grid and allowed to incubate for 20s, the grid is blotted from the back again for ~20s. The grid is then vitrified by plunging into liquid ethane and stored in liquid nitrogen. (B) SEM image of a grid after on-grid soaking of proteinase K crystals with I3C solution with hundreds of crystals to be potentially investigated. (C) High resolution SEM image showing crystal density and distribution. Scale bar 1mm in (B) and 50μm in (C).

### Identification and selection of ligand soaked crystals

Crystals on the TEM grid were identified in the SEM after platinum coating as described previously (Martynowycz et al., 2019; Martynowycz et al., 2019; Wolff et al., 2020). The crystal density was high with crystals being present over both the holey carbon film and the film above the grid bars (Figure 1). A target crystal was identified in both SEM and FIB imaging (Figure 2) next to two other crystals approximately at the center of a grid square. These crystals measured between 3-10μm across. The view of the back crystal was occluded in the FIB image at 18° by another crystal near the grid bar.

**Figure 2.**
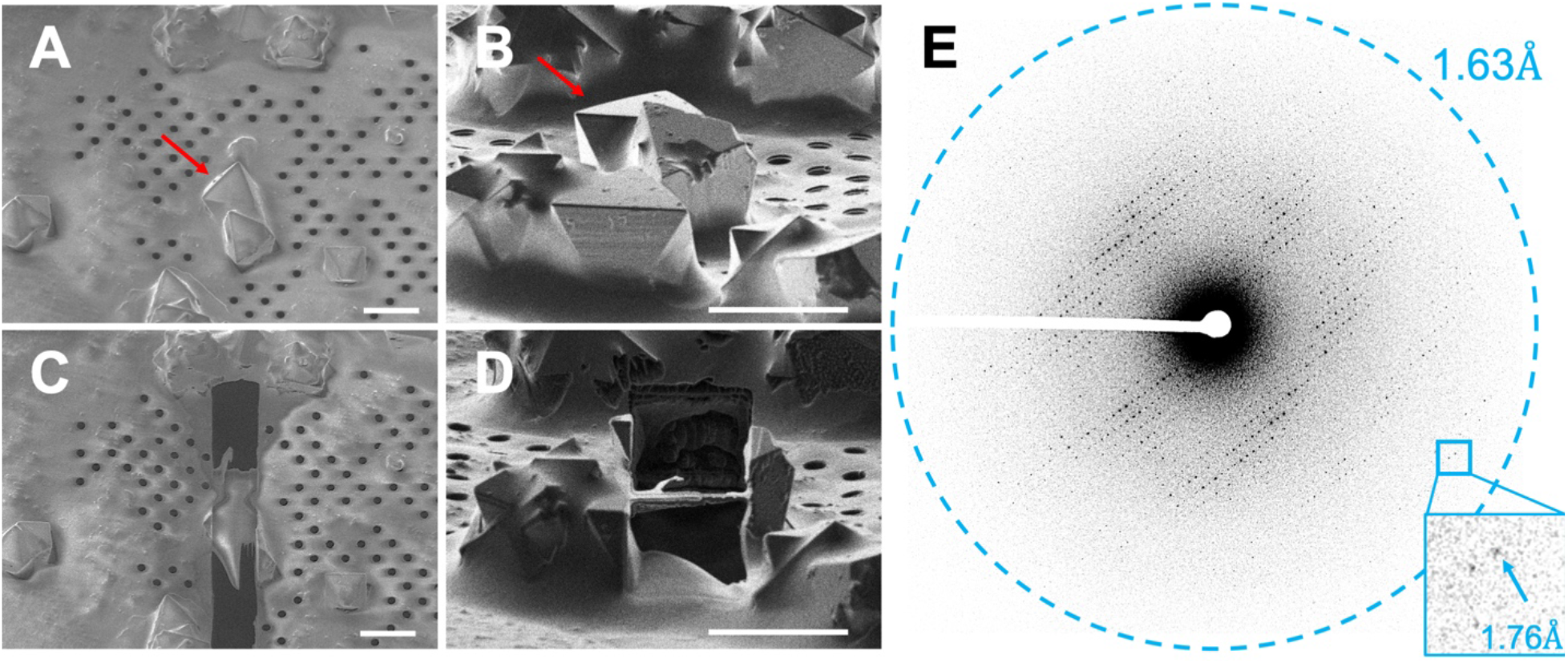
Cryo-FIB milling of a proteinase K microcrystal soaked with I3C. (A) SEM and (B) FIB images of a 10μm protein crystal before cryo-FIB milling. The crystal after milling the back crystal to a thickness of 200nm imaged using the (C) SEM and (D) FIB. (E) The resulting diffraction from this lamella at 0° for 1s. Inset showing visible high resolution spots. Scale bars 10μm in all images. Red arrow indicates crystal selected for investigation.

### Cryo-FIB milling

The occluding crystals in front of and abutting the crystal of interest were removed by milling to access the target crystal (Figure 2). Rough milling was done in cleaning cross sections from both the top and bottom of the desired lamella location. A final thickness of ~200nm was achieved after two polishing steps using a very low gallium beam current.

### MicroED data collection

The grid was cryo-transferred to a Talos Arctica TEM for MicroED data collection. Crystal lamellae were identified by taking a low-magnification montage within the MicroED software EPU-D (Thermo Fisher). The target lamella appeared as a long streak of white with a semitransparent lamella suspended in the gap. The crystal was screened for diffraction by collecting a single diffraction image at 0° for 1s using a 70μm selected area aperture to isolate the signal from a small area of the target crystal. Reflections to better than 2Å were observed (Figure 2) indicating that the target crystal was well ordered and suitable for MicroED data collection. A continuous rotation MicroED data set was collected over a total of 60°.

### MicroED data analysis

MicroED data were indexed, integrated, and scaled in DIALS as described (Clabbers et al., 2018; Parkhurst et al., 2016; Waterman et al., 2016; Winter et al., 2018). A resolution cutoff was applied after integration at 1.78Å, where the CC1/2 fell to a value of 0.33 (Table 1). This dataset from a single crystal was found to yield an overall completeness of 94.5%, a I/σI of 7.8, and R_pim_ of 14%.

**Table 1.** Refinement statistics for a proteinase K microcrystal soaked with I3C.

| **Proteinase K soaked on-grid with I3C** |  |
| --- | --- |
| Wavelength (Å) | 0.0251 |
| Resolution range (Å) | 43.32 - 1.78 (1.844 - 1.78) |
| Space group (#) | P 4<sub>3</sub> 2<sub>1</sub> 2 |
| Unit cell (a = b, c, α = β = γ) | 67.55, 102.77, 90 |
| Total reflections | 109326 (10458) |
| Multiplicity | 4.9 (4.8) |
| Completeness (%) | 94.52 (94.56) |
| Mean I/sigma(I) | 7.83 (2.19) |
| Wilson B-factor | 14.37 |
| R<sub>merge</sub> | 0.2622 (0.9953) |
| R<sub>pim</sub> | 0.1304 (0.5076) |
| CC<sub>1/2</sub> | 0.971 (0.365) |
| R<sub>work</sub> | 0.1694 |
| R<sub>free</sub> | 0.2094 |
| Number of non-hydrogen atoms | 2339 |
| macromolecules | 2029 |
| ligands | 64 |
| solvent | 246 |
| RMS (bonds) | 0.005 |
| RMS (angles) | 0.74 |
| Ramachandran favored (%) | 97.83 |
| Ramachandran allowed (%) | 2.17 |
| Ramachandran outliers (%) | 0 |
| Rotamer outliers (%) | 0 |
| Clashscore | 5.21 |
| Average B-factor | 13.51 |
| macromolecules | 12.59 |
| ligands | 19.44 |
| solvent | 19.61 |

### Structure determination and modeling

Molecular replacement (McCoy et al., 2007) was performed using the proteinase K model with PDBID 6CL7 to determine the structure. This model of proteinase K contains no solvent or ions in the model, making it ideal for molecular replacement when looking for ligands. A single, unambiguous solution in P 4_3_ 2_1_ 2 was found. The initial solution from molecular replacement was inspected for differences in the density corresponding to ligands. We found >25 peaks in the Fo-Fc map with values >6σ. From these peaks, four I3C ligands could be immediately identified in both the 2F_o_-F_c_ and F_o_-F_c_ maps (Figure 3). Prior to placing and refining the molecules, the rings of the I3C molecules were apparent - in some cases even without lowering the contour levels. The quality of these maps made for simple, unambiguous identification of the ligand molecules without any special treatment - such as Polder maps (Liebschner et al., 2017)-of the density.

**Figure 3.**
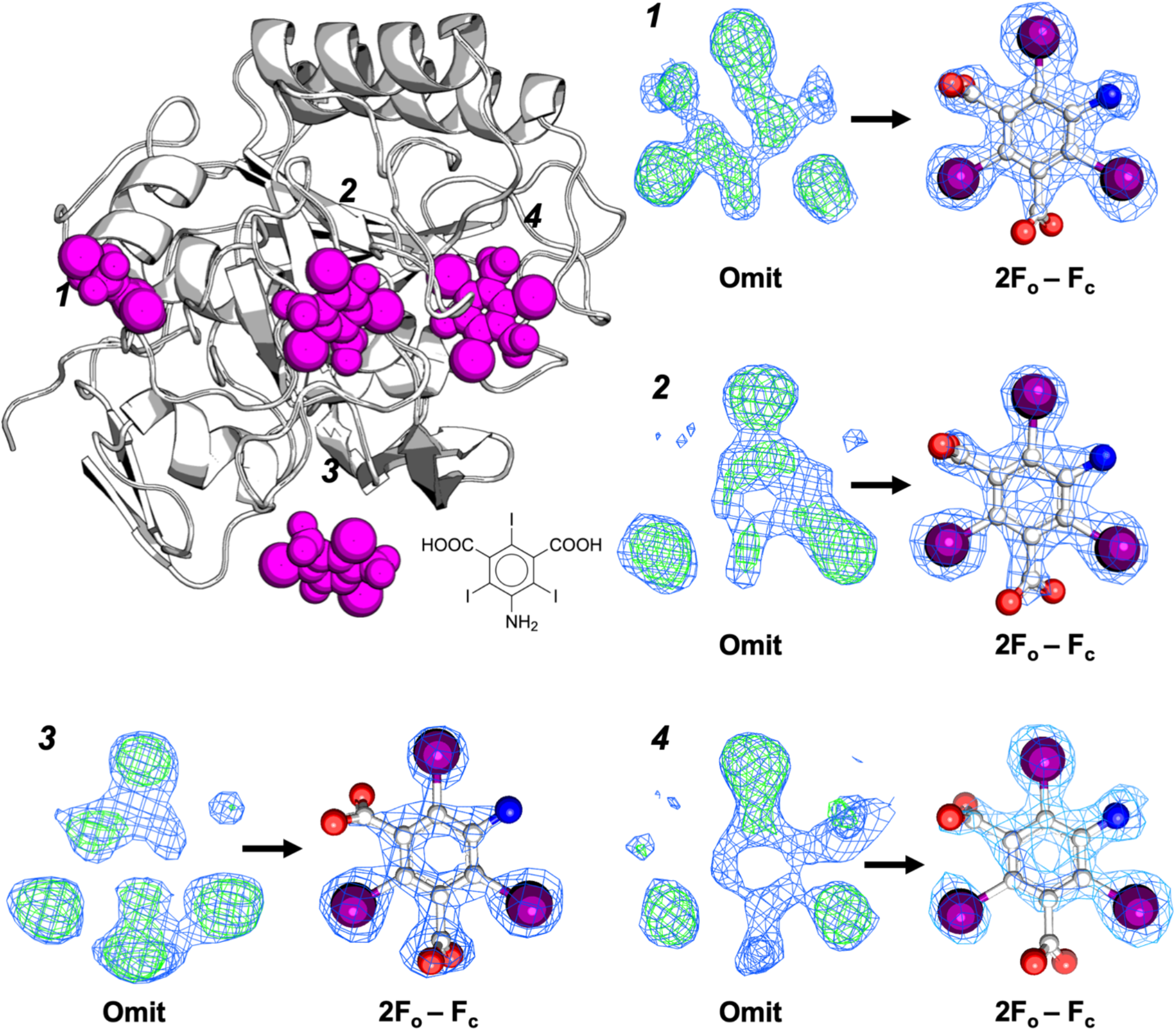
I3C ligands soaked into the protein lattice. (Top Left) Cartoon representation of the determined protein (beige) and the identified ligand binding sites in magenta spheres. Chemical structure of the I3C magic triangle appears at the bottom right of the protein cartoon. (Right and bottom) corresponding numbered sites with their densities before modeling (left side), and after the ligand was modeled and refined (right). Blue meshes are 2F_o_-F_c_ maps contoured at the 1.0σ level, and green meshes are F_o_-F_c_ maps contoured at the 3σ levels. The iodine atoms form an equilateral triangle with side lengths ~6Å.

### Structure refinement

After placing the four I3C ligands, the structure was refined and solvent molecules were added automatically using Phenix.refine (Afonine et al., 2012). Another round of refinement was conducted that allowed the occupancy of the I3C molecules to vary. The four I3C ligands were found to have occupancies between 58 and 75%, with average B-factors of 20 Å^2^. These B-factors are approximately equal to that of the modeled solvent atoms, and slightly higher than the average B-factor of the protein (13 Å^2^). The entire model was refined using individual B-factors, rather than group B-factors for the ligands, and resulting in a final R_work_ and R_free_ of 16.9% and 20.9%, respectively, attesting to the quality of the data. Detailed data collection and refinement statistics are presented in **Table 1**.

### Comparing soaking in X-ray crystallography with MicroED

Large proteinase K crystals were previously soaked with I3C and the structure determined by X-ray crystallography to 1.76 Å resolution with R_work_ of 14.0% and R_free_ of 19.1% (Beck et al., 2010). The X-ray study took advantage of anomalous scattering to identify the positions of the soaked I3C molecules and allowed the identification of 3 unique I3C molecules bound to the protein (Figure 4). Only one of the I3C ligands had high occupancy (75%) while the other two had occupancies of only 21% and 12%. In sharp contrast, soaking into much smaller crystals with MicroED identified 4 unique I3C molecules bound to Proteinase K. Importantly, all four I3C ligands have high occupancies – 75%, 61%, 59% and 58%.

**Figure 4.**
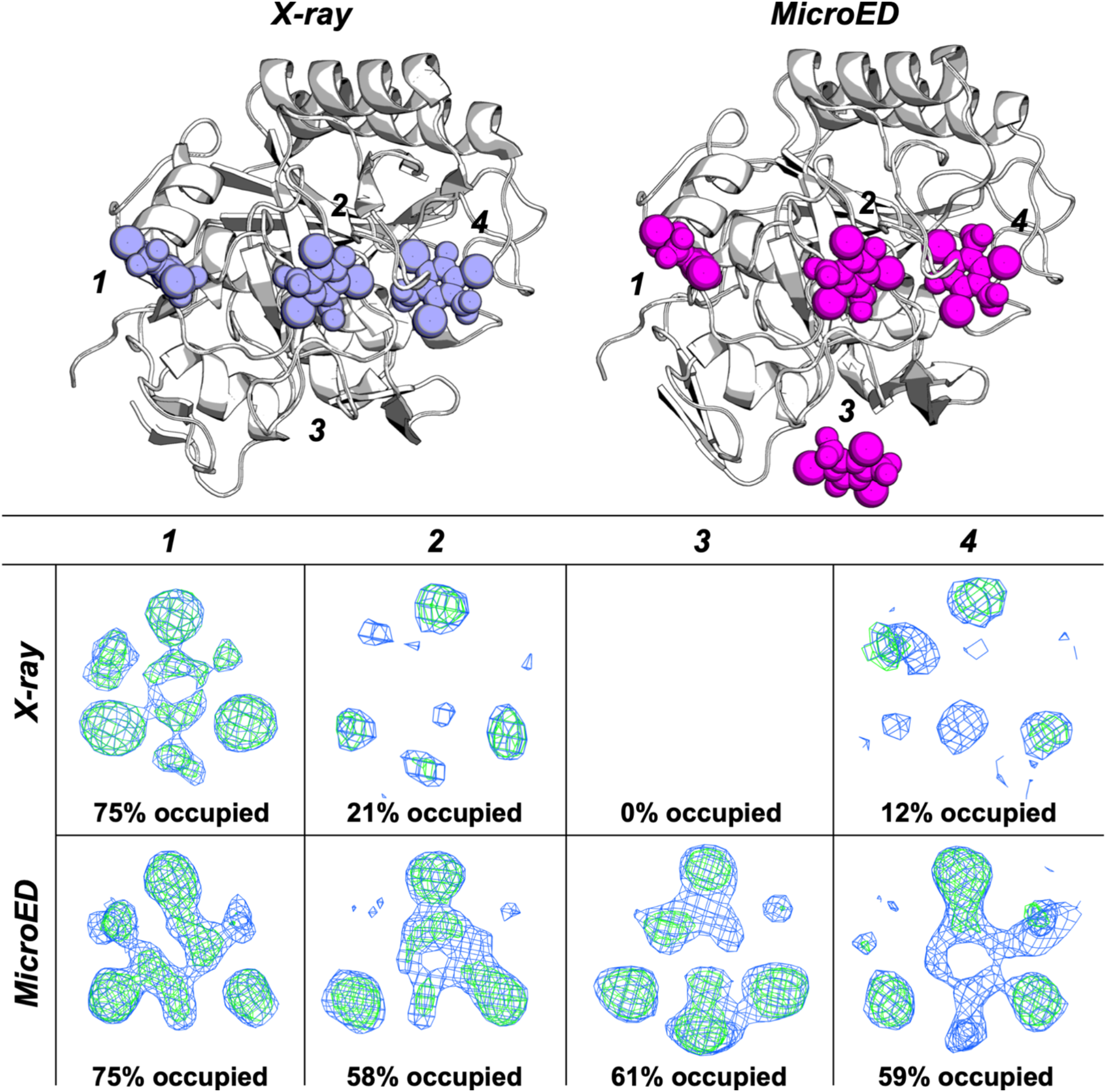
Comparison of ligand placement between a microcrystal determined by MicroED and a large crystal determined using X-ray diffraction. (Top left) MicroED (beige) and X-ray determined (light blue) structures overlaid with their bound ligands. The overall Ca RMSD between the models is 0.18 Å. (Right and bottom) For each numbered ligand, the corresponding 2F_o_-F_c_ and F_o_-F_c_ densities using only the protein phases are shown with the MicroED on the left and X-ray on the right for comparison. Ligand 3 in the X-ray structure was not observed. All 2F_o_-F_c_ maps are contoured at the 1s level, and F_o_-F_c_ maps are contoured at the 3s level. Maps are presented with a 1.75 Å carve for clarity. Density maps for each are calculated at 1.78 Å resolution. A sulfate ion appears in place of I3C in site 3 for the X-ray structure, and was omitted for clarity.

## Discussion

We demonstrate that soaking ligands into tiny crystals used for MicroED carries distinct advantaged over soaking experiments using large crystals for X-ray crystallography. Diffusion into small protein crystals is extremely efficient (Geremia et al., 2006). The Proteinase K soaking experiment with I3C presents an ideal test sample because a similar study was conducted by X-ray crystallography where both the resolution that was obtained (1.76 Å in X-ray and 1.78 Å in MicroED) and the refinement (Rwork of 14.0% and R_free_ of 19.1% in X-ray versus R_work_ and R_free_ of 16.9% and 20.9% in MicroED) were similar in both studies.

The crystals used for the X-ray study were at least 100x larger (Beck et al., 2010) than the crystals used here. Though the final resolutions were similar, our study identified an additional I3C ligand and all ligands had high occupancy and appeared clearer omit maps when compared with the X-ray study (Figure 4). The data suggest that ligands such as I3C may soak into much smaller crystals more efficiently without sacrificing map quality or resolution enabling rapid drug discovery pipelines to be utilized effectively.

Investigating ligand-protein interactions using on-grid soaking has several advantages to typical macromolecular X-ray crystallography ligand soaking pipelines. By placing an entire crystallization drop on the grid, hundreds or even thousands of micro-and nanocrystals can be soaked simultaneously in a single step. In X-ray crystallography traditionally crystals are individually looped, soaked into a high concentration ligand solution, back soaked into a cryo-protective solution, and then plunged into liquid nitrogen. This approach is lengthy, requires a high degree of dexterity and a lot of material and has multiple points of failure from operator error. The on-grid method presented here soaks thousands crystals at once, using less than 1% of the amount of ligand needed for X-ray crystallography, requires no manual manipulation of the individual crystals, and does not require a cryo-protective back soak, since crystals vitrified using supercooled ethane typically show no solvent rings (Figure 2). Moreover, the crystals used here are extremely small so they would pose a considerable challenge to accurately fish out with a nylon loop and optimally shoot at synchrotron end stations even with microfocus capabilities.

The additional step of a dual beam FIB/SEM to this method adds significant utility. Milling protein crystals has recently seen a high amount of interest for preparing crystals that are too small for synchrotron X-ray experiments, but still too large for MicroED. By milling the ligand soaked crystals we are no longer limited to crystals that happen to be in thin ice and happen to be smaller than 1μm thick, though this method is still tractable when a FIB/SEM is unavailable by means of crystal fragmentation (de la Cruz et al., 2017; Martynowycz et al., 2017). Using the FIB/SEM, even crystals large enough for synchrotron or home source X-ray experiments become amenable to MicroED.

On a single grid, crystals can be found in a variety of sizes (Figure 1). Since all the crystals on the grid have been subjected to the same soaking condition without differences between operators or soaking techniques, a systematic investigation of different crystal sizes and ligand occupancies is possible and would be of great interest. Furthermore, cryo-FIB milling of ligand soaked crystals in specific regions of individual crystals provides a unique opportunity for investigation of sub crystalline domains that may not be possible using other means.

We present a method for high throughput, on-grid soaking of ligands into protein microcrystals that is fast, simple, and effective. On-grid soaking of protein microcrystals is demonstrated and shown to effectively incorporate ligands. The results suggest that soaked microcrystals of the same protein-ligand complex have higher occupancy and number of ligands bound than using a much larger crystal required for X-ray experiments. Soaking microcrystals on TEM grids may even prove to be a preferable emerging method for synchrotron X-ray investigations. Recent results have shown that vitrified protein microcrystals on TEM grids can be located ahead of time using an SEM, and then subsequently used for X-ray diffraction experiments. Alternatively, rastering of the grid at synchrotron end points can also be used. We also envision applications for single particle cryoEM and cryotomography workflows where active compounds are added on-grid and allowed to interact just before vitrification, allowing for time resolved investigations of ligand activity on varying scales. The approach described herein has implications for drug discovery and optimization, the investigation of protein-ligand interactions, and investigating the properties of how ligands interact with and flow through protein crystals.

## Acknowledgements

This study was supported by the National Institutes of Health P41GM136508. The Gonen lab is supported by funds from the Howard Hughes Medical Institute. The structure factors and coordinates for the on-grid soaked Proteinase K structure will be deposited in the PDB. The associated maps will be deposited at the EMDB. We would like to thank Johan Hattne for useful discussions.

## Competing interests

A patent filing accompanies this work UCH-24160 UCLA GONEN 20200514 2020-883.

## Materials and Methods

### Materials

Proteinase K (*E. Album*) was purchased from Sigma and used without further purification. 5-amino-2,4,6-triiodoisophthalic acid (I3C) magic triangles were purchased from Hampton and prepared as described (Beck et al., 2010).

### Crystallization

Proteinase K crystals were grown in batch by dissolving 5mg of lyophilized proteinase K powder into 1 mL of 1.25M ammonium sulfate at 4°C. Crystals between 1-50μm were formed within one day, and typically microcrystals were visible under the light microscope within minutes. The crystal soaking solution was 1.25M ammonium sulfate 0.1M I3C.

### Grid preparation

Quantifoil R2/2 Cu200 grids were glow discharged for 30s immediately prior to use. Grids were loaded on to a Leica GP2 cryo-plunger inside of a cold room held at 4°C and 35% relative humidity. The blotting chamber was set to 4°C and 90% humidity. Filter paper was added prior to use, and the system was allowed to equilibrate for 15 mins prior to use. Grids were loaded into the plunger and 3μL of proteinase K slurry were applied to the carbon side (front) of the grid and allowed to incubate with the protein drop for 30s. The grid was then gently blotted from the back (copper side) for 20s. After blotting, 3μL of the crystal soaking solution (1.25M ammonium sulfate, 0.1M I3C) was added to the same grid and allowed to incubate again for 20s. The grids were then blotted again from the back for 20s and immediately plunged into liquid ethane and transferred to liquid nitrogen for storage.

### Cryo-FIB milling of the protein crystals

Stored grids were clipped with the carbon facing side up and transferred into a cryogenically cooled Thermo-Fisher Aquilos dual beam FIB-SEM for milling. Grid clips were marked with a dot on the top with a sharpie to indicate the milling direction. A thin layer of platinum (~10nm) was deposited on the grids by sputter coating prior to inspecting the grids using the SEM (Michael W Martynowycz et al., 2019). An additional layer (~300nm) of carbon-rich platinum was added on top of this using the gas injection system. We applied the GIS platinum layer in the mapping position to prevent shadowing of the platinum layer and opened the GIS valve at 12mm rather than 7mm in order to have a slower, more controlled deposition. Crystals were identified in the SEM and FIB, brought to eucentric height, and milled using cleaning cross sections in steps into lamellae until a thickness of ~200nm as described (Duyvesteyn et al., 2018; Michael W. Martynowycz et al., 2019). The final polishing step used a gallium beam current of 10pA to remove the last few nm of crystal from the lamellae (Michael W Martynowycz et al., 2019).

### MicroED data collection

MicroED data collection was performed very similarly to previous experiments (Hattne et al., 2019, 2018, 2015; Jason de la Cruz et al., 2019; Martynowycz et al., 2020; Michael W Martynowycz et al., 2019; Michael W. Martynowycz et al., 2019). Grids with milled crystals were transferred to a Thermo-Fisher Talos Arctica transmission electron microscope after rotating the grids by 90°. Rotation by 90° assures the tilt axis during MicroED data collection is perpendicular to the milling direction, allowing for a greater rotation range of the crystal lamellae. The TEM was operated at liquid nitrogen temperature at an accelerating voltage of 200kV. Data were collected between the real space wedge between −30 and +30° at a rate of 0.25 °/s. Frames were read out every 1s and binned by 2. The total exposure to the crystal lamellae was 2.4 e^-^Å^-2^. The diffraction distance was set to 1900mm which corresponds to a crystal to detector distance of 1853mm after taking post-column magnification into account. Camera length was calibrated using diffraction from an aluminum film prior to loading the protein grid and to check for any distortions in the diffraction (Clabbers et al., 2018, 2017).

### MicroED data processing

Data were converted from MRC to SMV format as described, and an ADSC offset of 512 was applied to compensate for negative bias and pixel truncation (Hattne et al., 2016). The data were indexed, integrated, and scaled in DIALS using the general linear background model using a frame pedestal of 512 and a detector gain of 14 (Clabbers et al., 2018; Parkhurst et al., 2016). A resolution cutoff of 1.78Å was selected by the CC_1/2_ = 0.33 criterion (Evans and Murshudov, 2013; Karplus and Diederichs, 2012). The structure was determined by molecular replacement in PHASER using the search model PDB 6CL7 using electron scattering factors (Hattne et al., 2018; McCoy et al., 2007). The TFZ and LLG of the solution were 69.1 and 6856, respectively. Four I3C molecules were visible in the density prior to any refinement or modeling. These are apparent as the I – I distance is known to be ~6Å (Beck et al., 2008). The four I3C molecules were placed manually in COOT using the get monomer command using the three letter code I3C (Emsley and Cowtan, 2004). The structure was refined in Phenix.refine using electron scattering factors after creating restraints in Elbow as described (Afonine et al., 2012; Moriarty et al., 2009). The final model was refined using individual isotropic B-factors for the protein, solvent molecules, and I3C molecules. Occupancy was allowed to vary for the I3C molecules. Omit maps for MicroED in Figure 3,4 were generated directly after molecular replacement without further modification. For comparison, the coordinates and structure factors from PDB 3gt4 were downloaded, all ligands and solvent were removed, and 2F_o_-F_c_ and F_o_-F_c_ maps were generated.

### Figures and tables

Figures were arranged in Microsoft Powerpoint. Images were adjusted and cropped in FIJI (Schindelin et al., 2012). Tables were arranged in Microsoft Excel. Protein models and meshes were generated using PyMol (Schrödinger LLC, 2014).

